# Engineering of *Salmonella* phages into novel antimicrobial Tailocins

**DOI:** 10.1101/2023.10.03.560654

**Authors:** Cedric Woudstra, Anders Nørgaard Sørensen, Lone Brøndsted

## Abstract

Due to the extensive use of antibiotics, the increase of infections caused by antibiotic resistant bacteria are now a global health concern. Phages have proven useful for treating bacterial infections and represent a promising alternative or complement to antibiotic treatment. Yet, other alternative exists, such as bacteria-produced non-replicative protein complexes that can kill their targeted bacteria by puncturing their membrane (Tailocins). To expand the repertoire of Tailocins available, we suggest a new approach transforming phages into Tailocins. Here we genetically engineered the virulent *Ackermannviridae* phage S117, as well as temperate phages Fels-1, -2 and Gifsy-1 and -2 targeting the food pathogen *Salmonella*, by deleting the *portal vertex* or *major capsid* gene using CRISPR-Cas9. We report the production of Tailocin particles from engineered virulent and temperate phages able to kill their native host. Our work represents a steppingstone to tape into the huge diversity of phages and transform them into versatile puncturing new antimicrobials.

## 1. Introduction

Bacteriophages (phages) are natural predators of bacteria that are concomitantly present wherever a suitable host are found. Phages have been used for more than one century to treat bacterial infection, mostly in Eastern countries (1). The Eliava Institute in Georgie for example (2), has been providing phage preparations to treat bacterial infections of *Staphylococcus, Streptococcus, Pseudomonas aeruginosa, Proteus, E. coli, Salmonella*, and others since 1923 (3). Today, phages are recognized as a possible alternative and/or complement to classical treatment (e.g. antibiotics) of bacterial infection and are successfully used to treat resistant bacterial infection (4,5). Due to the increase of acquired resistance to available antibiotic treatment, multidrug-resistant bacteria are currently responsible for about 15.5% of hospital acquired infection cases (6). One of the most spectacular and mediatic example is probably the story of Tom Patterson, who suffered an infection with *Acinetobacter baumannii*, and recovered miraculously after being injected with a personalized phage treatment (7).

While phages are promising antimicrobial agents for targeting antibiotic resistant bacteria, other strategies using phage-related particles can be devised. Tailocins, also called Bacteriocin phage-like particles, are natural non-replicative high molecular mass protein complexes that resemble phage tails with *Myoviridae* or *Siphoviridae* morphologies (8). Tailocins consist of a contractile (R-type) or non-contractile (F-type) sheath forming a rigid tube attached to a baseplate with a spike-shaped protein complex at its tip and associated receptor binding protein (RBP) (9). They kill by perforating bacteria cell wall and leading to membrane potential collapse (10). Tailocins have obtained increasing interest in the last decade for their potential use as therapeutics to fight antibiotic resistant bacteria (11,12). For example, a recent study characterized a Tailocin with a broad host range against *Burkholderia spp*., known to acquire antibiotic resistance and representing a significant threat to cystic fibrosis patients (13). Tailocins have also been used as biocontrol against phytopathogenic bacteria (14). For example, the Tailocin produced by *Pseudomonas fluorescens* SF4c was successfully used to control the bacterial-spot disease in tomatoes caused by *Xanthomonas vesicatoria* (15). Interestingly, Tailocins can be genetically engineered to change their host range by altering their RBPs. The RBP from the R2 Tailocin produced by *Pseudomonas aeruginosa* was replaced by the RBP from phage PS17, resulting in killing of a different subset of *Pseudomonas aeruginosa* strains than the native Tailocin (16). These few examples illustrate how Tailocins can be appealing as new antimicrobials. Moreover, Tailocins can theoretically kill one target cell per particle (R-type (10)), do not carry genetic material and therefore do not transduce DNA, and are non-replicative and therefore could be easier to validate administratively as protein complexes for therapeutic use (17).Yet, while Tailocins have been isolated in different bacteria genera, only a limited number are currently characterized (18).

To harness the therapeutic potential of Tailocins against bacteria causing human infections, we suggest a new approach engineering both virulent and temperate phages into versatile Tailocins by preventing head to tail connection. Considering the huge diversity of phages, transforming virulent or temperate phages into Tailocins may provide a large source of antimicrobials for targeting diverse pathogenic bacteria. In phage T4, the portal protein constituting the initiator complex of a 12-mer ring, and the major capsid protein that polymerises on the portal complex are required for correct head formation and head to tail assembly (19–21). Thus, the *portal* and *major capsid* genes constitute suitable target to delete to generate Tailocins from phage. As our bacterial target, we selected the food pathogen *Salmonella enterica* Typhimurium (*S*. Typhimurium), since clones showing antibiotic resistances are increasingly isolated (22). Among phages infecting *S*. Typhimurium, *Ackermannviridae* are attractive to develop into Tailocins as they encode up to five Tail Spike Proteins (TSPs) (23). Each individual TSP function as an RBP by recognizing specific O-antigen or K-antigen receptor from different bacterial hosts such as *Salmonella, Shigella, Enterobacter, Dickeya, Klebsiella and Serratia* (24). Thus, transformation of the broad host range *Ackermannviridae* phages into Tailocins may be used to fight O- and K-antigen expressing bacteria, such as *Salmonella* and *Klebsiella spp*. (25). Temperate phages Fels-1, -2, Gifsy-1 and -2 are widespread *Salmonella* temperate phages showing *Siphoviridae* and *Myoviridae* morphologies. While temperate phages Gifsy-1 and -2 bind to OmpC as receptors (26), Fels-1 and -2 receptors have not been experimentally determined but may be different from OmpC. Thus, genetic engineering of these temperate phages into Tailocins may transform *Salmonella* into a Tailocins factory producing a cocktail of contractile and non-contractile Tailocins particles targeting the same host but potentially different receptors.

In this study, we aimed to increase the versatility of phages by transforming them into Tailocins. *Ackermannviridae* virulent phage S117 as well as *S*. Typhimurium temperate phages Fels-1, -2, Gifsy-1 and -2, were genetically engineered using CRISPR-Cas9 to be deficient for the *portal* or the *major capsid* gene. Their deletion allowed production of Tailocin particles derived from phage S117 as well as Fels-1, -2, Gifsy-1 and -2 Tailocin particles that were characterized for their abilities to kill *S*. Typhimurium.

## 2. Materials and Methods

### 2.1. Bacteriophage, bacterial strain, and culturing media

*S*.Typhimurium LT2C devoid of prophages (27) was used as the host for engineering *Ackermannviridae* virulent phage S117 (24). *S*. Typhimurium 3674 was used as the native host for engineering temperate phages Fels-1, -2, Gifsy-1 and -2 (28). Luria agar 1,5% (LA), luria agar 0,6% and luria broth (LB) were used to cultivate *S*. Typhimurium on solid or in liquid media, with or without antibiotics (kanamycin 50 μg/ml, spectinomycin 50 μg/ml).

### 2.2. From phage DNA to cloned plasmids

Phage DNA was extracted using a rapid DNA extraction method (29). Briefly, a phage stock solution at 10^11^ pfu/ml was incubated with 50 µL DNase I 10x buffer, 1 µL DNase I (1 U/µL), and 1 µL RNase A (10 mg/mL) for 1.5 h at 37 °C without shaking. Following, 20 µL of 0.5 M EDTA (final concentration 20 mM) was added to inactivate DNase I and RNase A. The phage protein capsid was digested using 1.25 µL Proteinase K (20 mg/mL) and incubated for 1.5 h at 56 °C without shaking. DNA amplification of phage or bacterial genetic sequences were performed by PCR using the CloneAmp HiFi PCR premix® (Takara©). Primers used in this study are listed in Table S1. PCR amplicons were purified using the Zymo PCR purification kit® (Zymo Research©). Plasmid constructions were performed using In-fusion® HD-cloning kit (Takara©) in highly competent Stellar *E. coli* cells. Constructed plasmids were extracted using the GeneJET Plasmid Miniprep Kit® (Thermo Fisher Scientific©). All procedures followed the manufacturer’s instruction. DNA concentration was measured with Qubit (Thermo Fisher Scientific©) and DNA quality verified on a 1% agarose gel.Plasmids were sequence-verified by Sanger sequencing (Eurofins Genomics©).

### 2.3. Genetic engineering

A double CRISPR-Cas plasmids system (pEcCas, pEcgRNA (30)) was used to engineer phages S117, while phages Fels-1, -2, Gifsy-1 and -2 were engineered using homologous recombination linear DNA fragments. Plasmid pEcCas contains kanamycin resistance (Addgene Plasmid #73227) and plasmid pEcgRNA (Addgene Plasmid #166581) contains spectinomycin resistance. The plasmid pEcCas encodes Cas9 under a constitutive promoter, as well as the Lambda-Red system under an arabinose inducible promoter. The plasmid pEcgRNA contains the guide targeting Cas9 to a specific genetic region and the homologous recombination DNA sequence to delete the gene of interest. Importantly, strain *S*. Typhimurium LT2C showed to be not transformable with plasmid pEcCas, due to the presence of multiple restriction-modification (RM) systems. We overcame this problem by cloning the methylase *Stylti* (STM0357 from NC_003197.2) in *E. coli* and transformed pEcCas in our Stylti_m+ *E.coli*. We could then extract the pEcCas methylated in *E. coli* and transformed it back in *S*. Typhimurium LT2C.

### 2.4. CRISPR-Cas gRNA efficiency against S117

Each guide was evaluated for its efficiency against the phage S117. *S*. Typhimurium LT2C strain was transformed with pEcCas and pEcgRNA-g1 to pEcgRNA-g20. The protection provided by each guide was measured individually by comparing the reduction of infection by double layer plaque assay, when infected by serial dilution of phage S117. Briefly, LT2C strain containing pEcCas and pEcgRNA-g was grown exponentially until OD600 0.5. 100 µl of LT2C was mixed with 4 ml of top agar 0.6% and poured onto LA plate containing kanamycin (50 mg/ml) and spectinomycin (50 mg/ml) to select for bacteria containing both pEcCas and pEcgRNA. Serial dilutions of phage stock S117 were then spotted (10 µl) onto the surface of the dried plate. The plate was incubated at 37°C overnight and plaques were counted the next day and compared to a negative control of LT2C containing only the pEcCas plasmid spotted with the same serial dilution of S117. The guide showing the highest reduction in S117 infectivity was selected for further downstream application.

### 2.5. Construction of pEcgRNA plasmid to engineer S117

Primers used to construct pEcgRNA are available in Table S1. Plasmid pEcgRNA used to genetically engineer phage S117 was built in three steps. First, pEcgRNA was reverse amplified by PCR to include the most successful sgRNA guide under the constitutive promoter J23119, producing plasmid pEcgRNA-g. Second, plasmid pEcgRNA-g was reverse amplified by PCR and cloned together with the DNA homologous recombination (HR) template to allow knockout of the gene of interest, the *portal vertex* of S117, producing plasmid pEcgRNA-g-HR. The HR fragment contains two regions of 500 bp, the left homology arm (LHA) and the right homology arm (RHA) that flank the region to modify (here, the *portal* gene). LHA and RHA were amplified separately from phage S117 DNA and joined together by SOE PCR to form the HR recombination template (30). Third, pEcgRNA-g-HR was reverse amplified and cloned together with the complementation *portal* gene (*cportal*), producing the plasmid pEcgRNA-g-HR-c*portal* (Figure S1). The *cportal* gene was codon optimized for *S. enterica* to avoid pEcgRNA-g-HR-*cportal* being self-targeted. The *cportal* was ordered commercially (IDT®). The complementation *cportal* had 78% nucleotide similarity and 100% amino acid identity to phage S117 native portal protein. This allowed to produce fully functional phage particles containing S117 genome deleted for the *portal* gene.

### 2.6. S117 portal gene deletion

The LT2C strain containing pEcCas and pEcgRNA-g-HR-*cportal* was grown at 37°C until exponential phase at OD_600nm_ 0.5, together with arabinose (0.1%) to induce the Lambda-Red system on pEcCas. Then, 100 µl of culture was mixed with 4ml of LA 0.6% and poured onto LA plate containing kanamycin (50 μg/ml) and streptomycin (50 μg/ml). After drying for 10 min, 10 µl of serial dilutions of phage S117 stock (10^10^ pfu/ml) were spotted onto the surface and dried for 15 min. After overnight incubation, single plaques visible at the highest dilution were picked with toothpick and resuspended into SM phage buffer (31) and tested by PCR for the deletion of the *portal* gene. Plaques that were positive for the *portal* deletion were used in a new round of infection. The process was repeated three times to ensure the purity of the recombinant S117Δportal deleted for the *portal* gene.

### 2.7. Fels-1, -2, Gifsy-1, -2 major capsid gene deletion

*S*. Typhimurium strain 3674 containing pEcCas was grown at 37°C until exponential phase at OD_600nm_ 0.5, together with arabinose (0.1%) to induce the Lambda-Red system on pEcCas for homologous recombination. The cells were then harvested and washed 3 times with water. The cells were electroporated (MicroPulser Electroporator Bio Rad with settings: Ec1 (V=1.8 kV) for 0.1 cm cuvettes and Ec2 (V=2.5 kV) for 0.2 cm cuvettes) with 250 ng of linear PCR product of a homologous recombination fragment targeting Fels-1 major capsid protein gene (MCP) to replace it with a chloramphenicol cassette CmR (Figure S2). The electroporated cells were resuspended into 500 µl of warmed LB media and incubated for 1h at 37°C under shaking at 180 rpm. 100 µl, 10 µl, 1 µl were then spread onto LA containing kanamycin and chloramphenicol at 50 μg/ml and incubated overnight at 37°C. Single colonies were then tested by PCR to check for the deletion. A single colony containing pEcCas and deleted for Fels-1 *major capsid* gene was chosen. The clone empty of Fels-1 *MCP* was then used, and the procedure was repeated with a new homologous recombination PCR product targeting the *MCP* of Fels-2, then Gifsy-1 and then Gifsy-2, with a different antibiotic cassette (Figure S3, S4, S5), until producing a strain engineered to be devoid of all *major capsid* genes from Fels-1, -2, Gifsy-1 and Gifsy-2.

### 2.8. Tailocin production: S117 Tailocin

One litre of exponentially grown LT2C was infected with S117Δ*portal* at MOI (multiplicity of infection) 0.1, 1, 10 and 100. 35 ml samples were taken after 10, 20, 30, 40 and 50 minutes of infection to evaluate the phage adsorption efficiency and the optimal time for Tailocin particles release. Samples were then centrifuged at 8,000 rpm for 10 minutes at 4°C. The supernatant was then filtered twice with 0.35 µm and 0.22 µm filter and kept at 4°C for further use. Presence of phage and Tailocin particles were assessed by spot assay on *S*. Typhimurium LT2C. Tailocin particles were further concentrated by PEG precipitation. A solution of PEG_8000_ at 30% in a solution of 1,5M NaCl (autoclaved) was used to precipitate the Tailocin particles, by adding 1/3 of the PEG solution to 2/3 of the Tailocin particles. After 48h at 4°C, the suspension was centrifuged at 12,000 rpm for 2h, and the supernatant discarded. The pellet was then resuspended in 500 µl of SM phage buffer.

### 2.9. Tailocin production: Fels-1, -2, Gifsy-1, -2 Tailocins

One litre of exponentially grown strain 3674 deleted for the *major capsid* genes of Fels-1, -2, Gifsy-1, -2 was submitted to mitomycin C treatment at 2 µg/ml for 3h. The cells were then centrifuged for 20 minutes at 8000 rpm and the supernatant was sterilized using 0.45 µm and then 0.22 µm filters. The supernatant was kept at 4 °C for two weeks prior being used to allow for the degradation of the mitomycin C.

### 2.10. Tailocin killing assay

100 µl of exponentially grown *S*. Typhimurium strain was mixed with 4 ml of LA 0.6% and poured onto LA plate. The Tailocin solutions were then tested by spotting 10 µl of serial dilutions onto the bacterial lawn. A result was considered positive when a lysis ring was observed at the place of the drop.

## 3. Results

### 3.1. Overall approach for engineering of phage S117 into a Tailocin

We devised a two-step approach to produce Tailocin from phage S117. First, we engineered the genome of phage S117 by deleting the *portal* gene (S117Δ*portal*) using CRISPR-Cas9 during infection of *S*. Typhimurium LT2C carrying a complementation *portal* gene *in trans* (Figure 1a) This produced S117Δ*portal* phage particles that lack the *portal* on the genome but phenotypically behave as wild type phages. Second, we used the phage stock of S117Δ*portal* to infect the native host, *S*. Typhimurium LT2C, leading to production of Tailocin particles (Figure 1b). Thus, to engineer the genome of phage S117, we used a CRISPR-Cas9 approach already proven to efficiently produce phage mutants (32).

**Figure 1.**
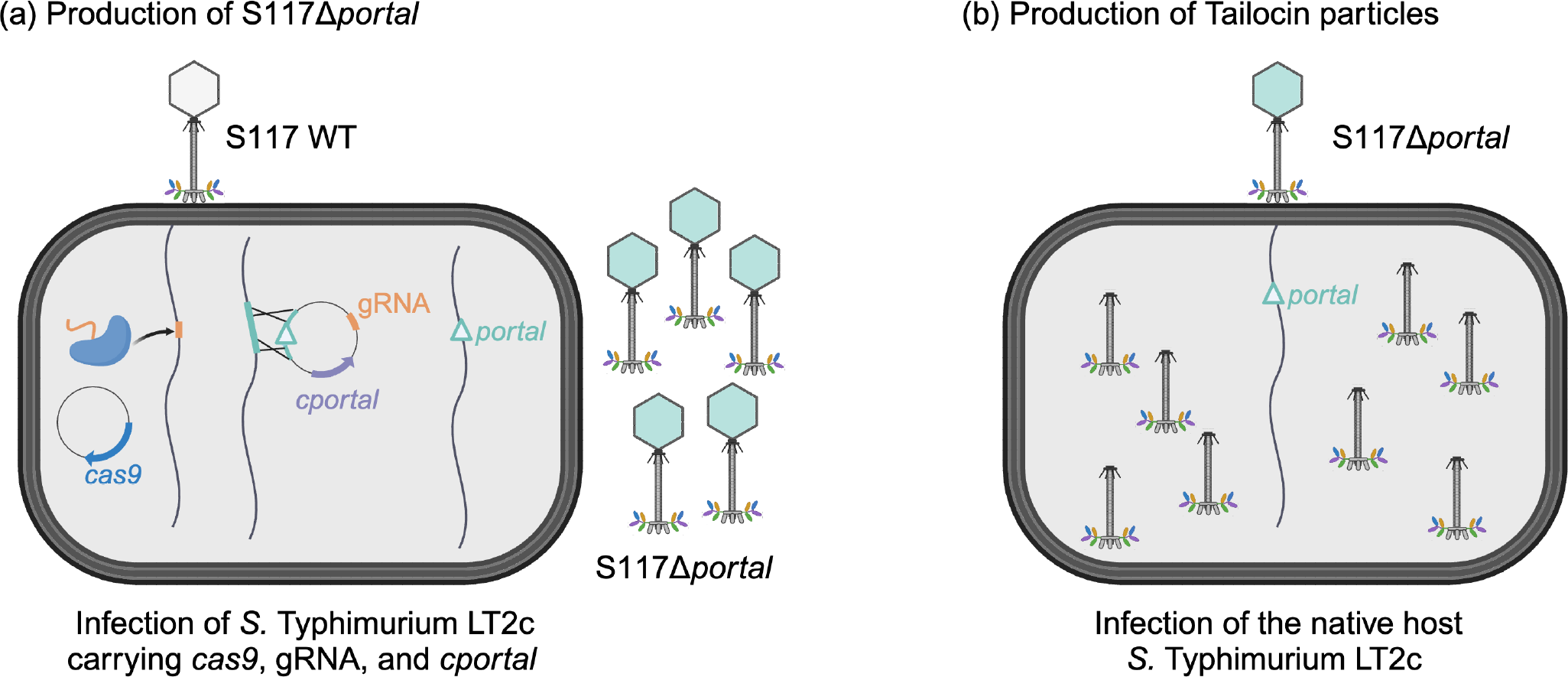
Engineering strategy to produce Tailocin particles from phage S117. **(a)** Production of phage S117D*portal*. A native phage S117 infects *S*. Typhimurium LT2C with plasmids pEcCas containing *cas9* and pEcgRNA. A guide RNA on the pEcgRNA plasmid allow Cas9 to target and cut the wild type phage genome at the *portal* gene. pEcgRNA contains the homologous recombination sequences allowing the phage genome to escape Cas9 by deleting the *portal* gene. In presence of the complementation *cportal*, the portal protein is produced constitutively and allows production of phage particles containing the phage genome deleted for the *portal* gene. **(b)** Production of Tailocin particles from phage S117D*portal*. The engineered phage phage S117D*portal* is used to infect the native S. Typhimurium LT2C host. In the absence of any *portal* gene to complement the engineered phage genome, the portal protein is not synthesized and only Tailocin particles are produced.

### 3.2. Engineering of phage S117 lacking the portal gene

#### 3.2.1. CRISPR-Cas9 guides efficiencies

A key aspect of using CRISPR is to provide the right guide for Cas9 to reach its target (33). Especially, modified phage DNA makes it difficult to predict for the guide efficiency (33). Phage FEC14 and CBA120, which also belong to the *Ackermannviridae* family, have already been shown to be resistant to enzymatic restriction (34,35), due to DNA modification from a deoxyuridylate hydro-methyltransferase. The phage S117 genome (genbank accession number MH370370) also contain a locus (*orf51* to *orf55*) predicted to be involved in the production of non-canonical nucleotides by substituting thymine by hydroxy-methyluracil.

Therefore, phage S117 is expected to show variability in the efficiency of different CRISPR-guides. Consequently, we evaluated 20 different guides targeting the *portal* gene of phage S117 for their efficiency to allow Cas9 to restrict the S117 genome in the *portal* gene. The *portal* gene of phage S117 (*orf149*) is located within the locus of head and tail assembly and guides were selected manually distributed over the entire 1683 bp gene and further checked bioinformatically using CrisprScan (36). The efficiency of each guide was calculated based on the reduction in the Efficiency Of Plating (EOP, (37)). Phage S117 plaque formation was assessed on a lawn of *S*. Typhimurium LT2C containing pEcCas9 and the guide to the *portal* from phage S117. The results were compared with the number of plaques formed on a lawn of LT2C without pEcCas9 (Table 1). As expected, we found that guides efficiency to reduce EOP of S117 was variable and did not correlate with software prediction, as previously reported for phage T4 (33). EOP reduction varied from no reduction up to five log reduction in S117 efficiency to infect LT2C. Based on the results, guide 11 showing a five-log reduction of EOP was selected as the more efficient to target S117 *portal* gene.

**Table 1.**
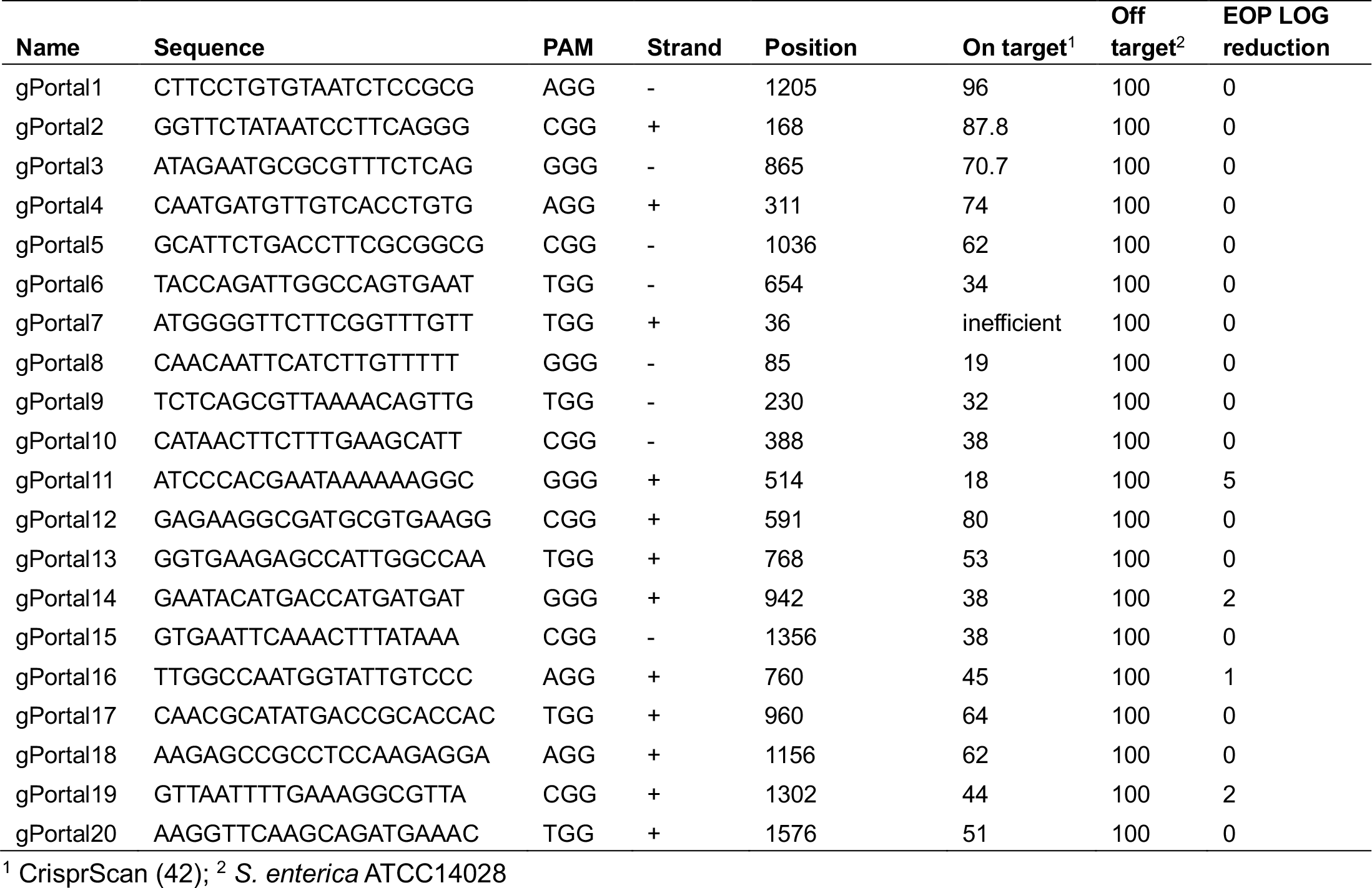
Predicted and tested efficiency of CRISPR-Cas9 guides targeting S117 *portal*.

#### 3.2.2. Production of engineered S117Δ*portal*

To produce phage S117 lacking the *portal* gene, we used wild type phage S117 to infect *S*. Typhimurium LT2C containing Cas9 (plasmid pEcCas), guide 11, the homologous recombination (HR) fragments LHA and RHA fragments surrounding the *portal* gene, and the complementation c*portal* gene *in trans* (cloned onto pECgRNA, Figure S1). LHA and RHA were selected to be 500bp long surrounding the *portal* gene to allow for the genetic recombination exchange to delete the *portal* gene. Single plaques were selected to test for the presence of the genetic recombination. PCR confirmed that the plaques corresponded to engineered S117Δ*portal* (Figure 2). Some plaques contained a mix of engineered S117Δ*portal* and native S117 (Figure 2). From the plaques containing S117Δ*portal*, three successive rounds of plaques purification and sequencing were performed to ensure only engineered phages S117 lacking the *portal* were isolated. In summary, after selection of the best CRISPR-Cas9 guide, it was possible to genetically engineer S117 to lack the *portal* gene.

**Figure 2.**
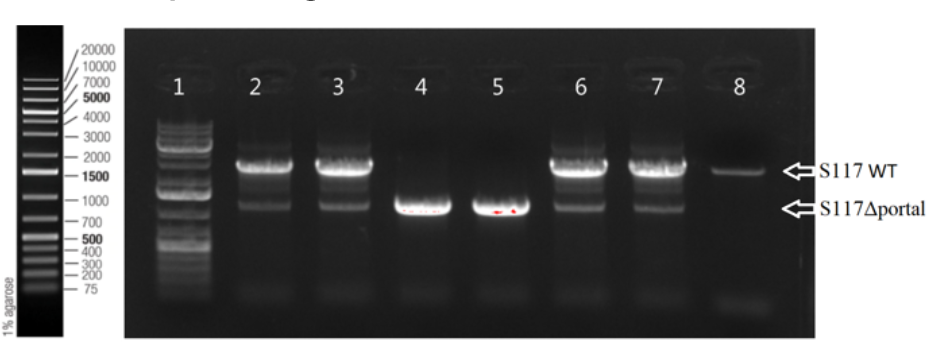
Plaque PCR of S117Δ*portal* to confirm the absence of the *portal* gene. Presence of the wild type *portal* gene results in a band of 3,688 bp (well 8, genomic DNA of S117), but if deleted the amplification produce a band of 1,191 bp. Plaques from well 2, 3, 6 and 7 contain a mix of engineered and native phages. Plaques from wells 4 and 5 contain engineered S117Δ*portal* only. Ladder of 1 Kb in well 1.

### 3.3. Characterization of the engineered phages S117Δportal

#### 3.3.1. S117Δ*portal* replication characterization

To characterize the engineered phage, S117Δ*portal* was propagated on *S*. Typhimurium LT2C containing the *cportal* complementation *in trans* (pECgRNA, Figure S1), producing a S117Δ*portal* phage stock of 3.2 x 10^11^ pfu/ml when plaquing on this strain (Figure 3a). In contrast, S117Δ*portal* could not form single plaques in the absence of the complemented *cportal* and only formed lysis zone up to 3 x 10^6^ pfu/ml on lawns of *S*. Typhimurium LT2C (Figure 3b). This confirmed that S117Δ*portal* was biologically incapable of producing functional phage particles in the absence of the *cportal* provided *in trans*.

**Figure 3.**
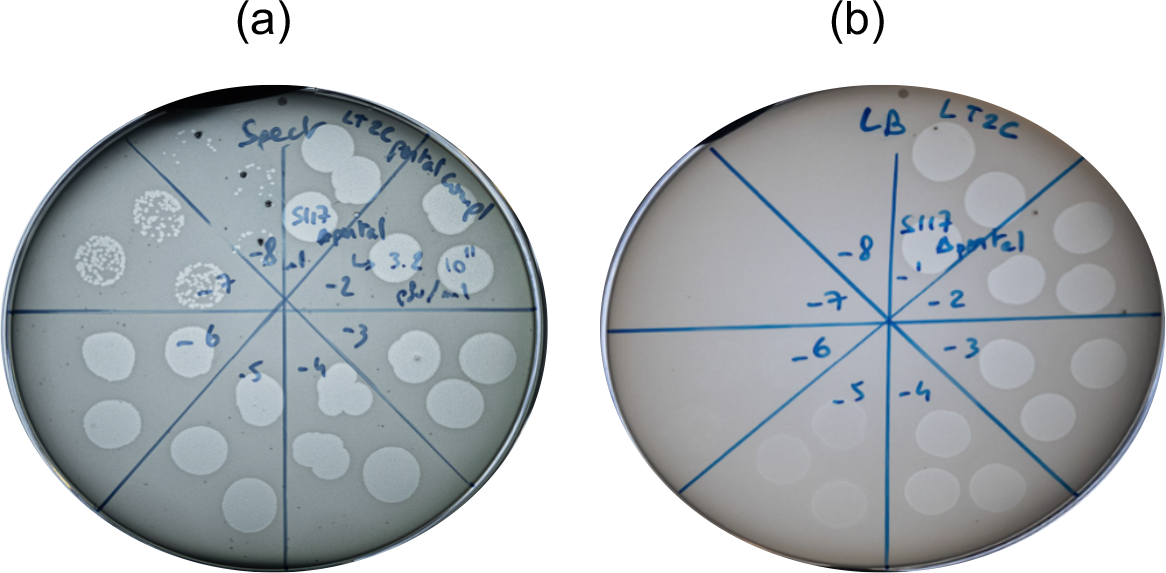
Biological confirmation of engineered S117 deleted for the *portal* gene. (a) Phage S117Δ*portal* infection on a lawn of *S*. Typhimurium LT2C containing the complementation *portal* gene. Single plaques are produced up to dilution 10^-7^ and 10^-8^. (b) Phage S117Δ*portal* infection on a lawn of *S*. Typhimurium LT2C without the complementation *portal* gene. No single plaques are visible.

#### 3.3.2. Production of S117Δ*portal* particles

We further investigated the production of phage particles from S117Δ*portal*. S. Typhimurium LT2C carrying the complementation *cportal* was infected by phage S117Δ*portal* at an MOI of 1. Production of phage particles as well as the effect on the host population were monitored (Figure 4a) by measuring phage free particles in the supernatant and the remaining host cells after spinning down.

**Figure 4.**
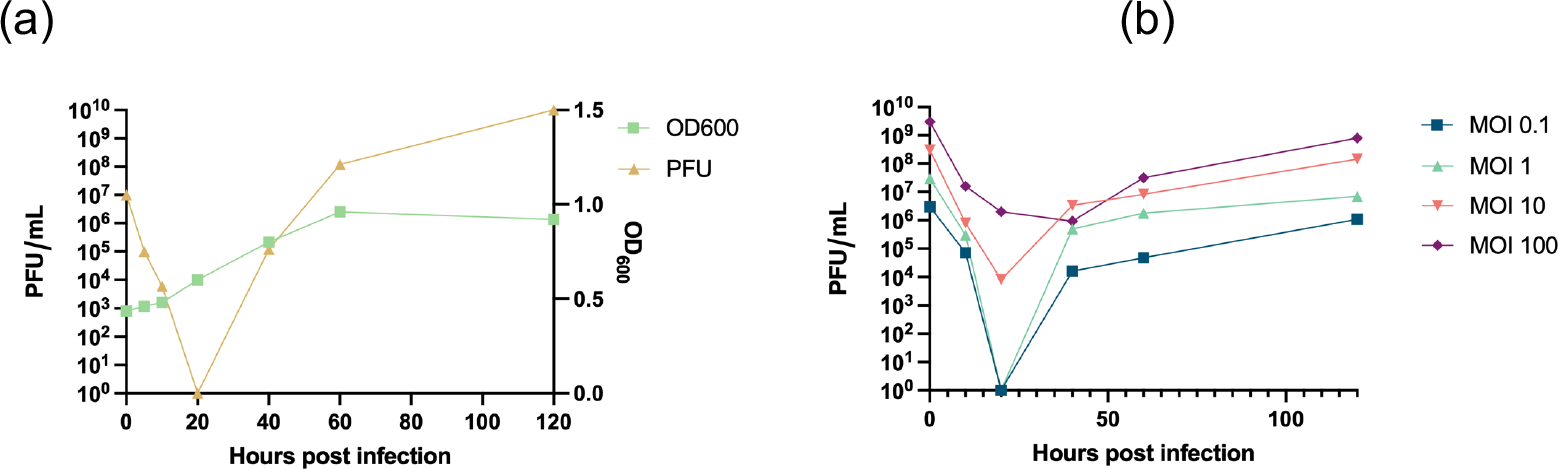
Determination of S117Δ*portal* phage particles release at different MOI. **(a)** Phage particles released at an MOI of 1. On the x axis are the time points in hours of the host culture. On the y axis on the left is the amount of phage calculated as plaque forming unit (pfu) per ml. On the y axis on the right are the OD_600_ values of *S*. Typhimurium LT2C carrying the complementation *portal* during infection. **(b)** Phage particles released at different MOI. On the x axis are the time points in hours of the host culture. On the y axis on the left is the amount of phage calculated as plaque forming unit (pfu).

At MOI 1, 100% of the phage particles of S117Δ*portal* absorbed onto their host within 20 minutes. The first release of new phage particles was observed at 40 minutes post infection and resulting in 10^6^ phage particles per ml. These results are concordant with the previous study of kuttervirus phage CBA120 also belonging to the *Ackermannviridae* family (34). The experiment was repeated with 0.1, 10, and 100 MOI to determine the influence of the MOI on the infection kinetics (Figure 4b). At an MOI of 0.1, all phage particles were adsorbed after 20 minutes post infection, whereas at MOIs 10 and 100, not all phage particles were adsorbed at 20 minutes post infection. After 40 minutes post infection, the amount of phage particles present was 10^4^, 10^6^ and 10^7^ pfu/ml for MOI 0.1, 10 and 100 respectively. Importantly, at MOI 10 and 100, the total amount of phage particles detected also included those that did not bind immediately to the host, whereas the phage particles found at MOI 0.1 and 1 only included those produced as there was no remaining free phage particles after 20 minutes post infection. Therefore, a lower MOI allows for a higher production of S117 phage particles at early post infection time points. In summary, we successfully constructed phage S117 lacking the *portal* gene, which when complemented by the *portal* protein *in trans* showed similar growth pattern as other *Kuttervirus* phages.

### 3.4. Tailocin particles produced from phage S117Δportal form inhibition zones

To produce Tailocin particles, it is important to notice that once the Tailocins are produced they must be harvested as soon as possible to avoid binding to a new host or cell debris and be irrecoverable. Therefore, we used the MOI 1 as the best compromise to produce the highest amount of Tailocin particles at 40 minutes post infection by infecting one litre culture of *S*. Typhimurium LT2C by S117Δ*portal*. The Tailocin particles produced were tested by spot assay onto the native hosts of phage S117 along with the native phage (24). The harvested Tailocin particles produced a lysis halo on all native S117 hosts and no clean lysis as when phage S117 was spotted (Figure 5a). Furthermore, no single plaques was observed, confirming the production of Tailocin particles and no phage particles. Tailocin particles were then concentrated using PEG (38) and a lysis halo and no single plaques were visible on a lawn of *S*. Typhimurium (Figure 5b). Moreover, the concentration of Tailocin was determined by mixing successive dilution of 10^8^ up to 10^1^ cfu of *S*. Typhimurium LT2C and counting the survival population. The estimated amount of Tailocin produced from one liter of culture and concentrated by PEG was 10^6^ particles per ml, considering one Tailocin kills one bacterial cell.

**Figure 5.**
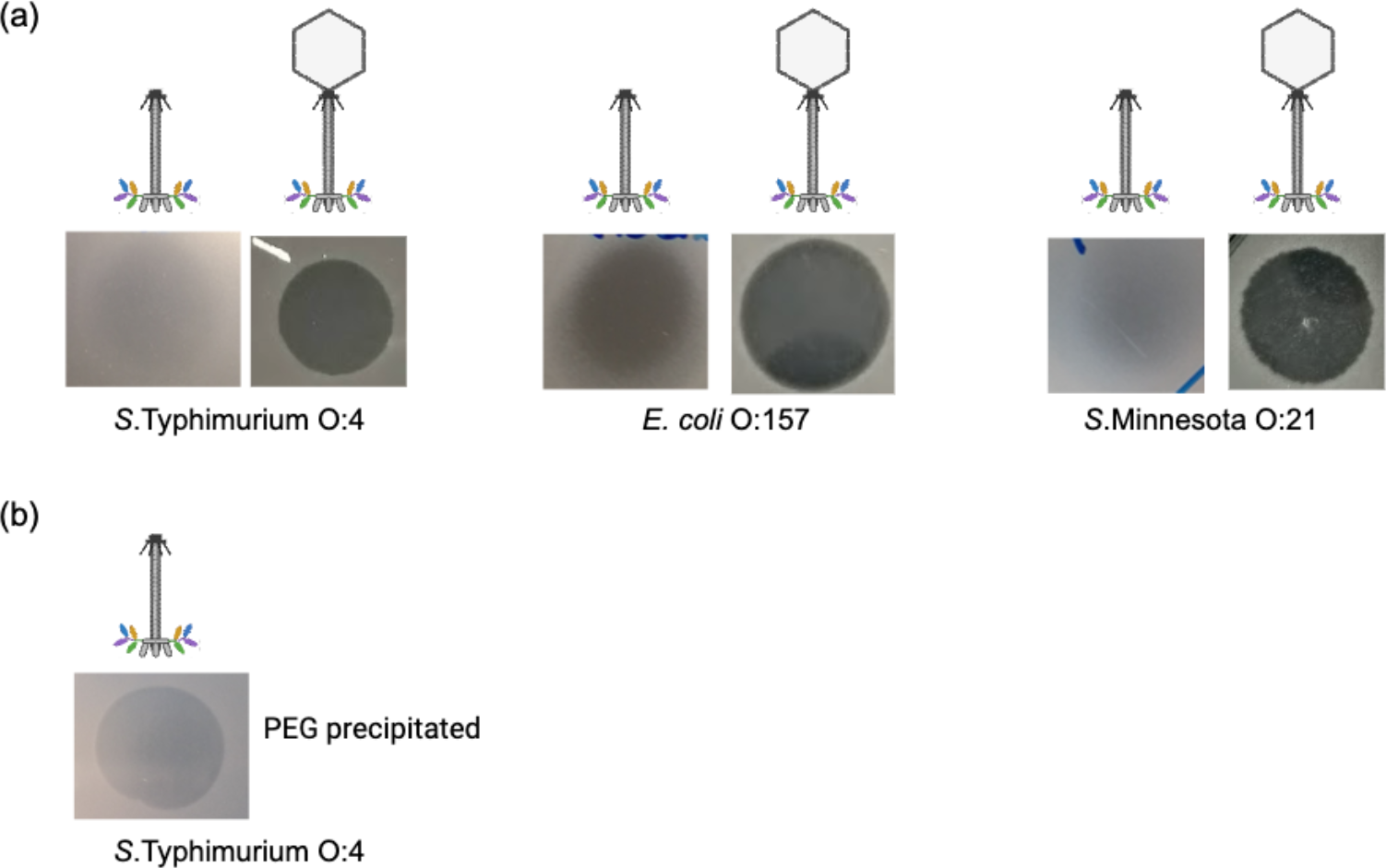
Effect of Tailocin on the native host of S117. **(a)** All three hosts *S*. Typhimuirum (strain LT2C (O:4)) and *S*. Minnesota (strain JEO2 (O:21)) and *E. coli* (strain NTCT12900 (O:157)) are sensitive to the Tailocin and the native phage S117. **(b)** Tailocin effect on *S*. Typhimurium after concentration 1000 times using PEG. A lysis halo is visible and no single plaques, confirming the presence of Tailocin particles

Visualization of the Tailocin particles were not attempted by TEM as a minimum of 10^9^ particles of phage are usually required (39). However, when serial dilutions of the concentrated Tailocin particles were spotted on a lawn of LT2C we noticed a few single plaques, suggesting the presence of phage particles at 10^2^ pfu per ml in the Tailocin preparation. Surprisingly, plaque PCR showed that these phage particles contained S117Δ*portal* and the pECgRNA plasmid carrying the *cportal* gene used for complementation. Therefore, the plaques are the result of a double infection by a transducing particle containing the *cportal* (pEcgRNA) and phage S117Δ*portal*, allowing S117Δ*portal* particles to be formed at a low frequency. We tested both our stock of S117Δ*portal* and S117 and found that both were able to transduce plasmids (pEcCas, pECgRNA) and chromosomal fragments from their host (data not shown). We estimated the presence of 10^4^ transductants per plaque-forming unit, in agreement with previous work (40). We then attempted to complement the *portal* on the chromosome to minimize the number of transducing particles containing the *cportal*, yet with little improvement (data not shown). Thus, we report the ability of this phage to transduce DNA, as previously described for other *Ackermannviridae* phages (40). Overall, we successfully produced Tailocin particles from phage S117.

### 3.5. *Tailocin particles produced by engineering temperate phages of S*. Typhimurium

Temperate phages may be easier to engineer as they are integrated in the host chromosomal genome. Seemingly as for virulent phage S117, the deletion of a structural head genes in all temperate phages would allow to produce a cocktail of different Tailocins at once. Temperate phages Fels-1, -2, Gifsy-1 and -2 are *Salmonella* temperate phages showing *Siphoviridae* and *Myoviridae* morphologies that could be transformed into Tailocins.

#### 3.5.1. Bioinformatic identification of Fels-1 and Fels-2, putative receptors

While temperate phage Gifsy-1 and -2 bind to the OmpC as receptors (26), Fels-1 and -2 receptors have not been identified. Considering the tail fiber protein of Fels-1 (STM0926 in LT2 genome NC_003197) is 93% and 92% identical to Gifsy-1 and Gifsy-2 respectively, we can assume Fels-1 use OmpC as a receptor. In contrast, Fels-2 encodes a tail fiber protein (STM2706 in LT2 genome NC_003197) with a C-terminal region similar to temperate phages SJ46 and ST64B, showing 84% and 72% identity, respectively. Since the tail fiber C-terminal part is responsible for binding to the bacterial receptor (41), we can assume Fels-2 would use the same receptor as phages SJ46 and ST64B. Unfortunately, neither SJ46 nor ST64B have their bacterial receptor characterized. Yet, as Fels-2 tail fiber is different from Fels-1 and Gifsy-1 and -2, we can assume Fels-2 uses a different receptor. Genetically engineering these prophages into Tailocins would therefore transform *Salmonella* into a Tailocins production factory producing a cocktail of contractile and non-contractile Tailocins particles targeting the same host but likely different receptors.

#### 3.5.2. Genetic engineering of temperate phages Fels-1, Fels-2, Gifsy-1, and Gifsy-2

To transform phages Fels-1, Fels-2, Gifsy-1, and Gifsy-2 into Tailocins, we successfully deleted their *major capsid* genes in *S*. Typhimurium 3674 by providing a PCR fragment containing a homologous recombination template deleted for the *major capsid* gene (Figure S2, S3, S4 and S5). Deletion of the *major capsid* genes were confirmed by PCR (figure 6a). We then induced the production of Tailocin particles upon activation of the temperate phages through mitomycin C treatment and concentrated it using PEG, as previously described for S117Δ*portal*. We tested for the presence of Tailocin particles by spot assay on a lawn of *S*. Typhimurium strain LT2C and 3674. Lysis was observed only on a lawn of LT2C (Figure 6b) but not on 3674. As *S*. Typhimurium LT2C is devoid of prophages, but not strain 3674, this may indicate a mechanism of superinfection exclusion. The amount of Tailocin particles produced was comparable to the amount obtained by engineering phage S117 (10^6^ particles per ml). It is probable that the low quantity of Tailocin particles obtained is in relation with the temperate phages’ RBPs targeting their own producer host. As previously observed for S117 Tailocin particles, due to the low quantity of particles obtained from temperate phages we could not visualize the Tailocin by TEM pictures. In summary, we successfully produced Tailocin particles from temperate phage present in *S*. Typhimurium.

**Figure 6.**
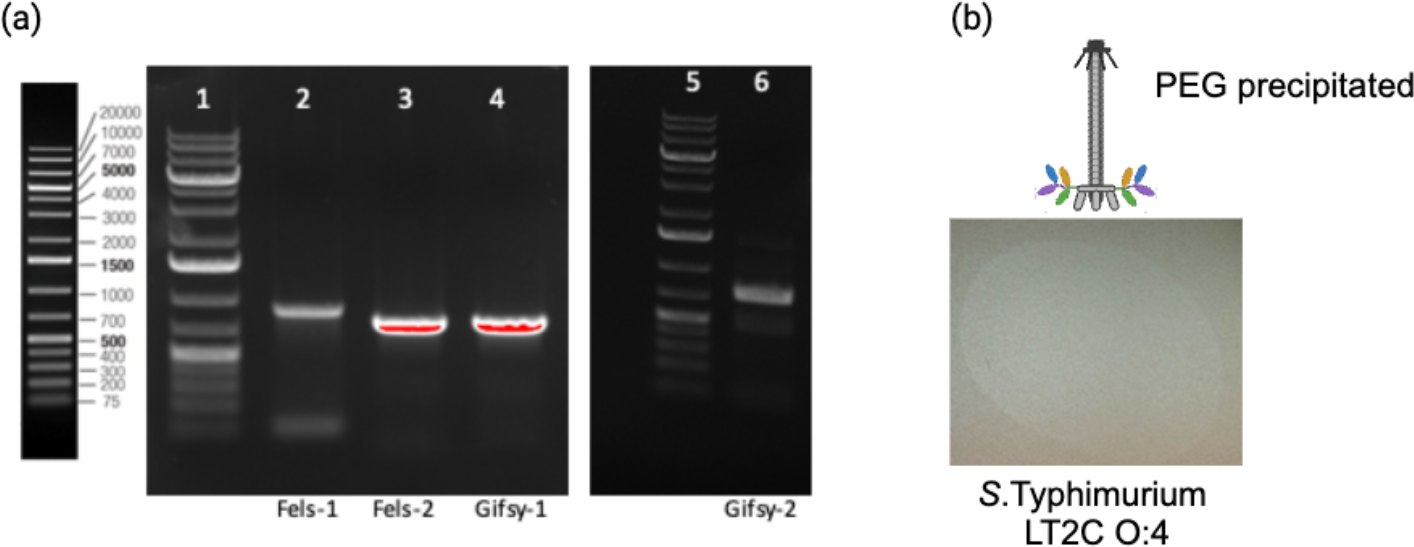
Construction and characterization of Tailocins originating from temperate phages Fels-1, Fels-2, Gifsy-1, and Gifsy-2 . **(a)** Deletion of major capsid genes from Fels-1, -2, Gifsy-1, -2 in S. enterica 3674. Wells 1) and 5) Ladder 1Kb+, 2) Fels-1, 3) Fels-2, 4) Gifsy-1, 6) Gifsy-2. An amplification can only be the result of the integration of the HR fragment into the chromosome, as the forward primer binds to the chromosome outside the HR and the reverse primer binds to the HR fragment inside the resistance gene inserted. All temperate phages are deleted for their major capsid. **(b)** Effect of Tailocin particles produced from temperate phages Fels-1, -2, Gifsy-1, -2 deleted for the major capsid gene on *S*. Typhimurium strain LT2C. A lysis halo is visible and no single plaques, confirming the presence of Tailocin particles produced from the engineered temperate phages of *S*. Typhimurium 3674.

## 4. Discussion

Phages have been used to successfully treat bacterial infection since a century in Eastern countries (e.g. Georgia (4)). Today, phages are starting to be accepted as viable complement / alternative to antibiotic treatment in Western countries too, especially to treat antibiotic resistant bacterial infection (42). Yet, due to their replicative nature, phages are difficult to validate pharmokinetically as well as setting up administrative schemes and are used mainly for compassionate treatments (42). Interestingly, bacteria can also produce phage-like particles that they use to kill competing species (43). These are called Tailocins and resemble head-less phage particles (44). Being inspired by the natural Tailocins, we proposed that phages may be transformed into such bacterial killing agents, by preventing head formation or head to tail connection of phage assembly. In this study, we genetically engineered virulent and temperate phages into Tailocins, by deleting the *portal* or the *major capsid* genes using CRISPR-Cas9 and homologous recombination, respectively.

Our engineering approach opens an avenue for transforming any type of phage into Tailocins. As a supplement, other approaches to produce Tailocin from phage particles may be considered. We previously attempted to produce Tailocin from phage S117 by osmotic shock and chemical treatment, without success (45), yet other phages may be transformed into Tailocins by these methods. Other genetic strategies to produce Tailocin exist, such as the use of RNA inhibition (46). An antisense RNA produced constitutively within the host, prior to infection by the native phage, could prevent the head synthesis by complementary binding. We have attempted this approach by targeting the *major capsid* gene of S117, without success (data not shown). In addition, CRISPR-Cas can also be used to prevent RNA synthesis through CRISPR-Cas inhibition (CRISPRi) (47), where Cas9 is engineered to be deficient for its catalytic site (deadCas9), preventing its endonuclease activity, yet preserving its binding ability (48). By targeting the promoter of the *portal* or the *major capsid* gene with the dCas9, RNA polymerase may be prevented from producing the mRNAs of these genes. For phage S117 this approach was not successful (data not shown), most likely due to few possibilities for guide design at the promoter region. In addition, the high level of DNA modification of the phage genome may prevent dCas9 to bind efficiently to the guide, similar to our many guides designed targeting the *portal* gene. Thus, these approaches may not be suitable for phages carrying modified DNA and deleting the *portal* or the *major capsid* genes using CRISPR-Cas and homologous recombination may be a more feasible way forward.

For therapeutic applications, the concentration of Tailocin particles is of major importance. Using our experimental setup, a production of up to 10^6^ particles per ml, after a concentration step, was achieved for both engineered virulent and temperate phages. Still, the quantity of particles produced in our study was lower than what was obtained for Pyocin production (49). This native Tailocin is encoded by the chromosome of *Pseudomonas* and its production is thus comparable to Tailocins derived from temperate phages. However, previous work reported that Pyocin particles may bind and/or contract upon encountering cellular debris (16). Thus, for improving production, we propose to change the RBP of the engineered temperate phage to prevent Tailocin binding to the producer host. This may result in upscaling the concentration of Tailocin particles up to a similar level as for Pyocin, where 10^11^ particles per ml of lysate can be obtained (16). This may be sufficient for therapeutical application as the recommended phage density is 10^8^ per ml (50). Yet, considering the replicative nature of phages, it is expected that the concentration of Tailocin particles needed is higher. Indeed, it has been reported that 3.10^11^ particles of Pyocin R2 injected intra peritoneal rescued a mouse from a *Pseudomonas aeruginosa* infection (49). This stress the importance of a substantial number of Tailocin particles for therapeutic applications.

Considering the huge number of phages present in the environment, we expect that many diverse phages can be transformed into Tailocins to target any pathogenic or unwanted bacteria. For selecting phages to transform into Tailocins, both target bacterium, morphology of the phage as well as the level of identified biological functions may be important. In our case *Akermannviridae* phage S117 was selected due to its contractile tail, characterized genome and multiple TSPs targeting diverse bacteria including the food borne pathogen *Salmonella* (24). Importantly, we successfully transformed this phage into a functional Tailocin preparation killing *Salmonella* by constructing an engineered phage S117D*portal* and propagating this on the native host. Yet, after concentrating the Tailocin, a low number of transducing particles were observed in the preparation. Since transducing particles are able to transfer genetic material horizontally, such particles are undesirable in products for therapeutic applications. To overcome this, non-transducing phages could be used as a backbone for engineering. Alternatively, switching Tailocin production to a cell-free expression system may be the future. We have tried to produce full phage particles and Tailocins from phage S117 and our genetically engineered S117Δ*portal*, in a cell-free system without success, yet the genome of phage S117 is quite large. However, cell-free expression systems have previously been used successfully to produce therapeutic amount of phage particles (51,52), yet our work highlight the need for optimization for large genomes as phage S117 (53). Overall, cell free extracts is an attractive strategy as it relies only on the genetic material provided. Thus, when further optimised, it could be envisioned to produce Tailocin particles out of any phages deleted for their *portal* or *major capsid* gene.

In summary, by optimizing engineering strategies as well as production, Tailocins are representing an immense untapped reservoir of alternative antimicrobials that could be use in the fight against antibiotics resistant bacteria.

## 5. Conclusions

With the increase of antibiotic resistant infections worldwide, the need for new alternatives has never been more important. Tailocins are promising antimicrobials that have already been tested *in vivo* and proved successful. Our study showed that it is possible to engineer Tailocins from phages, both virulent and temperate. We expect this work to provide a starting point for extending the range of Tailocin applications and new antimicrobials to treat bacterial infection in the future.

## Supplementary Materials

Figure S1: pEcgRNA plasmid map to delete S117 portal gene; Figure S2: HR PCR fragment to delete Fels-1 in strain *S*. Typhimurium LT2C; Figure S3: HR PCR fragment to delete Fels-2 in strain *S*. Typhimurium LT2C; Figure S4: HR PCR fragment to delete Gifsy-1 in strain *S*. Typhimurium LT2C; Figure S5: HR PCR fragment to delete Gifsy-2 in strain *S*. Typhimurium LT2C. Table S1: Primers used in this study.

## Author Contributions

Conceptualization, C.W. and L.B; Methodology, C.W.; Validation, C.W.; Formal analysis, C.W.; Investigation, C.W. and A.N.S.; Writing – original draft, C.W.; Writing – review & editing. C.W., A.N.S. and L.B.; Project administration, L.B.; Supervision, L.B; Visualization, C.W., A.N.S, and L.B; Project administration, L.B.; Funding acquisition, L.B. All authors have agreed to the submitted version of the manuscript.

## Institutional Review Board Statement

Not applicable.

## Informed Consent Statement

Not applicable.

## Data Availability Statement

Not applicable.

## References

1. Hatfull GF, Dedrick RM, Schooley RT. Phage Therapy for Antibiotic-Resistant Bacterial Infections. Vol. 73, Annual Review of Medicine. 2022.

2. Chanishvili N, Myelnikov D, Blauvelt TK. Professor Giorgi Eliava and the Eliava Institute of Bacteriophage. Vol. 3, PHAGE: Therapy, Applications, and Research. 2022.

3. Kutateladze M, Adamia R. Phage therapy experience at the Eliava Institute. Med Mal Infect. 2008;38(8).

4. Terwilliger A, Clark J, Karris M, Hernandez-Santos H, Green S, Aslam S, et al. Phage therapy related microbial succession associated with successful clinical outcome for a recurrent urinary tract infection. Viruses. 2021;13(10).

5. Rodriguez JM, Woodworth BA, Horne BA, Fackler J, Brownstein MJ. Case Report: successful use of phage therapy in refractory MRSA chronic rhinosinusitis. International Journal of Infectious Diseases. 2022;121.

6. Murray CJ, Ikuta KS, Sharara F, Swetschinski L, Robles Aguilar G, Gray A, et al. Global burden of bacterial antimicrobial resistance in 2019: a systematic analysis. The Lancet. 2022;399(10325).

7. Zolnikov TR. Global Health in Action Against a Superbug. Am J Public Health. 2019;109(4).

8. Ghequire MGK, De Mot R. The Tailocin Tale: Peeling off Phage Tails. Vol. 23, Trends in Microbiology. 2015.

9. Michel-Briand Y, Baysse C. The pyocins of Pseudomonas aeruginosa. Vol. 84, Biochimie. 2002.

10. Kageyama M, Ikeda K, Egami F. Studies of a pyocin: III. Biological properties of the pyocin. J Biochem. 1964;55(1).

11. Bali V, Panesar PS, Bera MB, Kennedy JF. Bacteriocins: Recent Trends and Potential Applications. Crit Rev Food Sci Nutr. 2016;56(5).

12. Behrens HM, Six A, Walker D, Kleanthous C. The therapeutic potential of bacteriocins as protein antibiotics. Vol. 1, Emerging Topics in Life Sciences. 2017.

13. Yao GW, Duarte I, Le TT, Carmody L, LiPuma JJ, Young R, et al. A broadhostrange tailocin from Burkholderia cenocepacia. Appl Environ Microbiol. 2017;83(10).

14. Grinter R, Milner J, Walker D. Ferredoxin containing bacteriocins suggest a novel mechanism of iron uptake in Pectobacterium spp. PloS One. 2012;7(3).

15. Príncipe A, Fernandez M, Torasso M, Godino A, Fischer S. Effectiveness of tailocins produced by Pseudomonas fluorescens SF4c in controlling the bacterial-spot disease in tomatoes caused by Xanthomonas vesicatoria. Microbiol Res. 2018;212–213.

16. Williams SR, Gebhart D, Martin DW, Scholl D. Retargeting R-type pyocins to generate novel bactericidal protein complexes. Appl Environ Microbiol. 2008;74(12).

17. Benítez-Chao DF, León-Buitimea A, Lerma-Escalera JA, Morones-Ramírez JR. Bacteriocins: An Overview of Antimicrobial, Toxicity, and Biosafety Assessment by in vivo Models. Vol. 12, Frontiers in Microbiology. 2021.

18. Patz S, Becker Y, Richert-Pöggeler KR, Berger B, Ruppel S, Huson DH, et al. Phage tail-like particles are versatile bacterial nanomachines – A mini-review. Vol. 19, Journal of Advanced Research. 2019.

19. Quinten TA, Kuhn A. Membrane Interaction of the Portal Protein gp20 of Bacteriophage T4. J Virol.2012;86(20).

20. Arisaka F. Assembly and infection process of bacteriophage T4. Chaos. 2005;15(4).

21. Leiman PG, Kanamaru S, Mesyanzhinov V V., Arisaka F, Rossmann MG. Structure and morphogenesis of bacteriophage T4. Vol. 60, Cellular and Molecular Life Sciences. 2003.

22. The European Union Summary Report on Antimicrobial Resistance in zoonotic and indicator bacteria from humans, animals and food in 2019–2020. EFSA Journal. 2022;20(3).

23. Volozhantsev N V., Verevkin V V., Krasilnikova VM, Kislichkina AA, Popova A V. Complete Genome Sequence of Klebsiella pneumoniae Bacteriophage KpS110, Encoding Five Tail-Associated Proteins with Putative Polysaccharide Depolymerase Domains. Microbiol Resour Announc. 2023;12(5).

24. Sørensen AN, Woudstra C, Sørensen MCH, Brøndsted L. Subtypes of tail spike proteins predicts the host range of Ackermannviridae phages. Comput Struct Biotechnol J. 2021;19.

25. Herridge WP, Shibu P, O’Shea J, Brook TC, Hoyles L. Bacteriophages of Klebsiella spp., their diversity and potential therapeutic uses. Vol. 69, Journal of Medical Microbiology. 2020.

26. Ho TD, Slauch JM. OmpC is the receptor for gifsy-1 and gifsy-2 bacteriophages of salmonella. J Bacteriol. 2001;183(4).

27. Shin H, Lee JH, Kim H, Choi Y, Heu S, Ryu S. Receptor diversity and host interaction of bacteriophages infecting Salmonella enterica Serovar Typhimurium. PloS One. 2012;7(8).

28. Garcia-Russell N, Elrod B, Dominguez K. Stress-induced prophage DNA replication in Salmonella enterica serovar Typhimurium. Infection, Genetics and Evolution. 2009;9(5).

29. Jakočiūnė D, Moodley A. A rapid bacteriophage dna extraction method. Methods Protoc. 2018;1(3).

30. Li Q, Sun B, Chen J, Zhang Y, Jiang Y, Yang S. A modified pCas/pTargetF system for CRISPR-Cas9-assisted genome editing in Escherichia coli. Acta Biochim Biophys Sin (Shanghai). 2021;53(5).

31. Fortier LC, Moineau S. Phage production and maintenance of stocks, including expected stock lifetimes. Methods Mol Biol. 2009;501.

32. Hoshiga F, Yoshizaki K, Takao N, Miyanaga K, Tanji Y. Modification of T2 phage infectivity toward Escherichia coli O157:H7 via using CRISPR/Cas9. FEMS Microbiol Lett. 2019;366(4).

33. Duong MM, Carmody CM, Ma Q, Peters JE, Nugen SR. Optimization of T4 phage engineering via CRISPR/Cas9. Sci Rep. 2020;10(1).

34. Kutter EM, Skutt-Kakaria K, Blasdel B, El-Shibiny A, Castano A, Bryan D, et al. Characterization of a Vil-like phage specific to Escherichia coli O157:H7. Virol J. 2011;8.

35. Fan C, Tie D, Sun Y, Jiang J, Huang H, Gong Y, et al. Characterization and Genomic Analysis of Escherichia coli O157:H7 Bacteriophage FEC14, a New Member of Genus Kuttervirus. Curr Microbiol. 2021;78(1).

36. Moreno-Mateos MA, Vejnar CE, Beaudoin JD, Fernandez JP, Mis EK, Khokha MK, et al. CRISPRscan: Designing highly efficient sgRNAs for CRISPR-Cas9 targeting in vivo. Nat Methods. 2015;12(10).

37. Kutter E. Phage host range and efficiency of plating. Methods Mol Biol. 2009;501.

38. Yamamoto KR, Alberts BM, Benzinger R, Lawhorne L, Treiber G. Rapid bacteriophage sedimentation in the presence of polyethylene glycol and its application to large-scale virus purification. Virology. 1970;40(3).

39. Morgan G, Lim D, Wong P, Tamboline B. Electron microscopy to visualize T4 bacteriophage interactions with Escherichia coli strain DFB1655, an isogenic derivative of strain MG1655 engineered to express O16 antigen. Undergraduate Journal of Experimental Microbiology and Immunology (UJEMI). 2019;24(September).

40. Matilla MA, Fang X, Salmond GPC. Viunalikeviruses are environmentally common agents of horizontal gene transfer in pathogens and biocontrol bacteria. ISME Journal. 2014;8(10).

41. Islam MZ, Fokine A, Mahalingam M, Zhang Z, Garcia-Doval C, Van Raaij MJ, et al. Molecular Anatomy of the Receptor Binding Module of a Bacteriophage Long Tail Fiber. PloS Pathog. 2019;15(12).

42. Verbeken G, Pirnay JP. European regulatory aspects of phage therapy: magistral phage preparations. Vol. 52, Current Opinion in Virology. 2022.

43. Nolan LM, Allsopp LP. Antimicrobial Weapons of Pseudomonas aeruginosa. In: Advances in Experimental Medicine and Biology. 2022.

44. Scholl D. Phage Tail-Like Bacteriocins. 2017; Available from: 10.1146/annurev-virology

45. Woudstra C, Brøndsted L. Producing Tailocins from Phages Using Osmotic Shock and Benzalkonium Chloride. PHAGE. 2023;4(3).

46. Walker SA, Klaenhammer TR. An explosive antisense RNA strategy for inhibition of a lactococcal bacteriophage. Appl Environ Microbiol. 2000;66(1).

47. Netter Z, Boyd CM, Silvas T V., Seed KD. A phage satellite tunes inducing phage gene expression using a domesticated endonuclease to balance inhibition and virion hijacking. Nucleic Acids Res. 2021;49(8).

48. Saifaldeen M, Al-Ansari DE, Ramotar D, Aouida M. CRISPR FokI Dead Cas9 System: Principles and Applications in Genome Engineering. Vol. 9, Cells. 2020.

49. Scholl D, Martin DW. Antibacterial efficacy of R-type pyocins towards Pseudomonas aeruginosa in a murine peritonitis model. Antimicrob Agents Chemother. 2008;52(5).

50. Abedon ST. Phage therapy dosing: The problem(s) with multiplicity of infection (MOI). Bacteriophage. 2016;6(3).

51. Emslander Q, Vogele K, Braun P, Stender J, Willy C, Joppich M, et al. Cell-free production of personalized therapeutic phages targeting multidrug-resistant bacteria. Cell Chem Biol. 2022 Sep 15;29(9):1434–1445.e7.

52. Garenne D, Bowden S, Noireaux V. Cell-free expression and synthesis of viruses and bacteriophages: applications to medicine and nanotechnology. Vol. 28, Current Opinion in Systems Biology. 2021.

53. Rustad M, Eastlund A, Jardine P, Noireaux V. Cell-free TXTL synthesis of infectious bacteriophage T4 in a single test tube reaction. Synth Biol. 2018;3(1).

