## Supplementary information Figures S1-S5 for "Engineering of *Salmonella* phages into novel antimicrobial Tailocins"

Supplemental figures.

Figure S1. pEcgRNA plasmid map to delete S117 portal gene. Extracted from Snapgene.

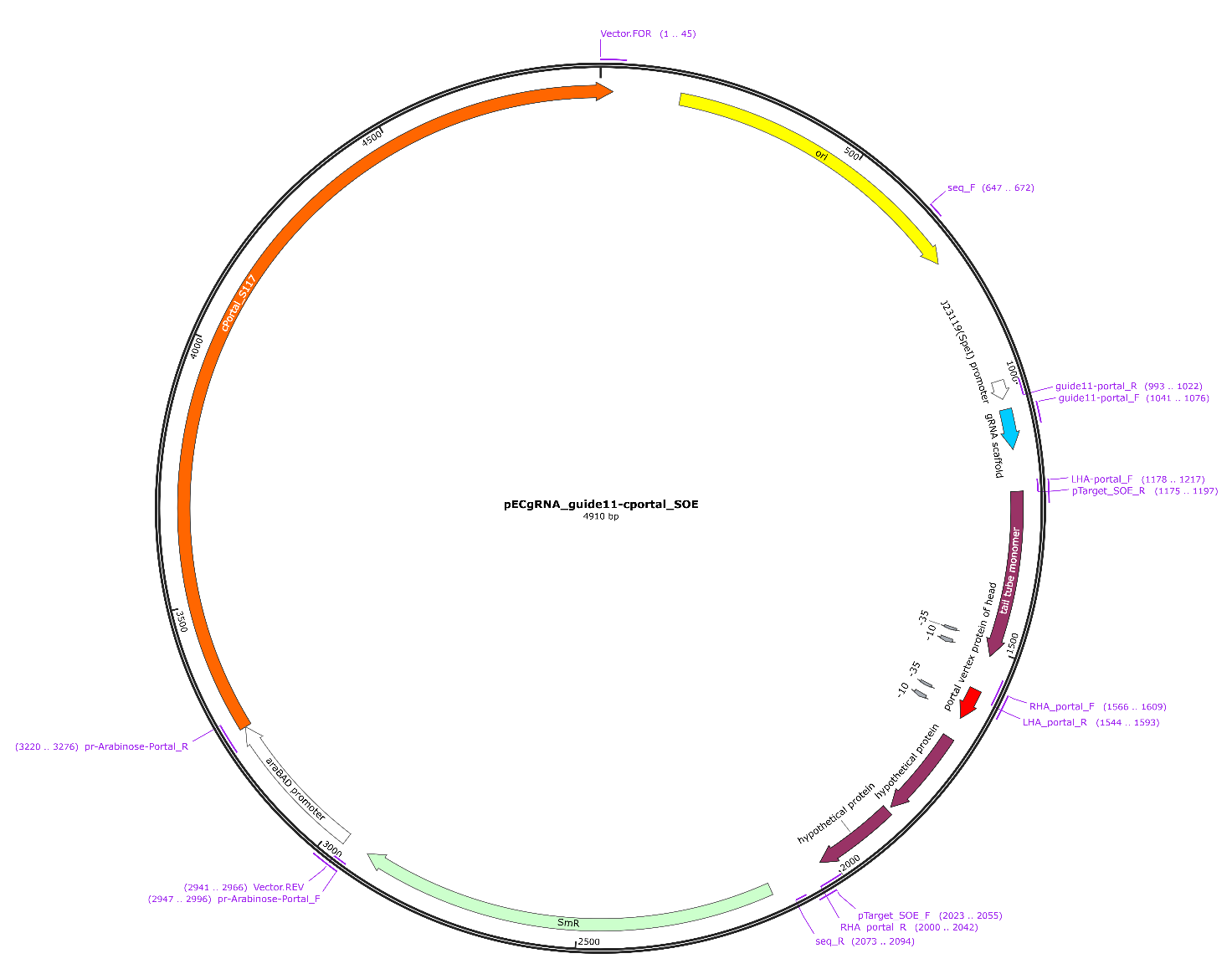

Figure S2. HR PCR fragment to delete Fels-1 in strain *S.* Typhimurium LT2C. Extracted from Snapgene.

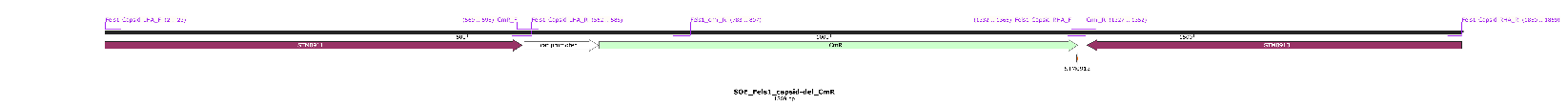

Figure S3. HR PCR fragment to delete Fels-2 in strain *S.* Typhimurium LT2C. Extracted from Snapgene.

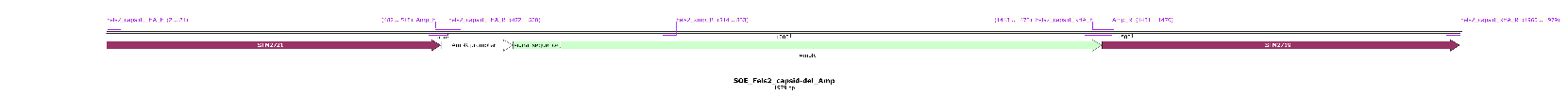

Figure S4. HR PCR fragment to delete Gifsy-1 in strain *S.* Typhimurium LT2C. Extracted from Snapgene.

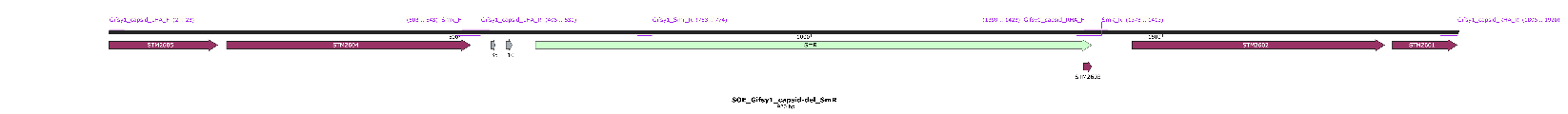

Figure S5. HR PCR fragment to delete Gifsy-2 in strain *S.* Typhimurium LT2C. Extracted from Snapgene.

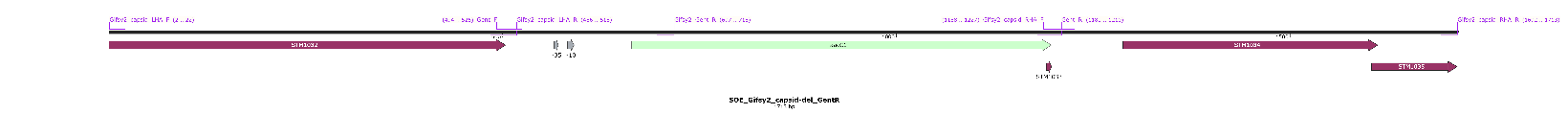
